# Latent dynamics of primary sensory cortical population activity is structured by fluctuations in the local field potential

**DOI:** 10.1101/2022.04.21.489039

**Authors:** Audrey Sederberg, Aurélie Pala, Garrett B Stanley

**Author notes:** Correspondence: Audrey Sederberg, School of Psychology and School of Physics Georgia Institute of Technology, Atlanta, GA 30332.

## Abstract

As emerging technologies enable measurement of precise details of the activity within microcircuits at ever-increasing scales, there is a growing need to identify the salient features and patterns within the neural populations that represent physiologically and behaviorally relevant aspects of the network. Accumulating evidence from recordings of large neural populations suggests that neural population activity frequently exhibits relatively low-dimensional structure, with a small number of variables explaining a substantial fraction of the structure of the activity. While such structure has been observed across the brain, it is not known how reduced-dimension representations of neural population activity relate to classical metrics of “brain state,” typically described in terms of fluctuations in the local field potential (LFP), single-cell activity, and behavioral metrics. Here, we relate the latent dynamics of spiking activity of populations of neurons in the whisker area of primary somatosensory cortex of awake mice to classic measurements of cortical state in S1. We found that a hidden Markov model fit the population spiking data well with a relatively small number of states, and that putative inhibitory neurons played an outsize role in determining the latent state dynamics. Spiking states inferred from the model were more informative of the cortical state than a direct readout of the spiking activity of single neurons or of the population. Further, the spiking states predicted both the trial-by-trial variability in sensory responses and one aspect of behavior, whisking activity. Our results show how classical measurements of brain state relate to neural population spiking dynamics at the scale of the microcircuit and provide an approach for quantitative mapping of brain state dynamics across brain areas.

**Author Summary:** Brain states have long been known to strongly shape sensory perception, decision making, cognition, and movement. Brain state during wakefulness changes constantly, classically assessed using changes in the spectral features of the local field potential (LFP) and behavioral measures. However, the connection between these classical measurements of brain state and the collective dynamics of populations of neurons is unclear. Here we fit a latent-variable model to population spiking activity, finding that latent variables inferred under the model are highly predictive of cortical state changes and that the latent dynamics are profoundly shaped by inhibitory cell activity. Our approach connects the activity patterns of ensembles of neurons to a classical measurement of brain state and opens new avenues for investigating brain state dynamics across diverse cortical areas.

## Introduction

Since the earliest electrophysiological measurements, it has been widely acknowledged that the brain undergoes gross changes in activity across a wide range of conditions related to sleep, arousal, attention, behavior, etc., collectively referred to as reflecting “brain state.” These changes are qualitatively apparent in measures such as the local field potential (LFP) and electroencephalogram (EEG) and have been conventionally quantified through changes in the spectral content across distinct frequency bands (1–5). Indeed, these measures have been shown to be strongly correlated with externally observable behavior, and thus capture, to some extent, behaviorally relevant changes in intrinsic brain activity. Although it has been shown in numerous studies that these measures of brain state are predictive of general levels of excitability, it remains unclear how they relate to the spiking activity of neural populations.

Descriptions of cortical state derive from decades of observations of behavior and careful circuit dissections of fluctuations in neural activity and behavior (6). Multiple studies have shown fluctuations in metrics of cortical activity are related to fluctuations in cognitive and sensorimotor behaviors in humans (7,8) as well as in animal models (9–13). A key model system for such studies is the mouse somatosensory cortex, in which states of wakefulness have been extensively characterized using the LFP and the activity patterns of single neurons (5,14–21). In the somatosensory cortex, “state” changes in spontaneous activity may be described on a continuum, from “deactivated” to “activated” (22), with the deactivated state characterized by local field potential (LFP) activity dominated by low-frequency components (1-10 Hz). The cortical state prior to sensory stimulation is predictive of trial-by-trial variation in the sensory response (16,23), and LFP state is informative of behavioral metrics, including whisking activity (5). Studies of single neurons have revealed cell-type-specific changes in firing rates with fluctuations in activation level (15,24,25), suggesting that the conventional notion of cortical state may be directly reflected in correlated fluctuations across the neuronal population, but this has not been studied in detail.

A picture that is emerging from analysis of simultaneous recordings of large populations of neurons is that, in many systems, a small number of latent variables explain much of the spatiotemporal structure of neural activity (26–29), and that frequently these latent variables are related to observed behavior (28,29). Analytic approaches based on inferring these latent dynamical variables can predict future neural activity as well as improve decoder performance in brain-machine interface applications (30). However, the connection between inferred latent variables and established measures of brain state is not well understood.

Here, we use unsupervised learning to model spontaneous spiking activity recorded in mouse somatosensory cortex during wakefulness and relate the population spiking states inferred from the model to classical LFP-based measures of state in mouse S1. We ‘open up the black box’ of the model to examine the structure of learned states, finding that putative inhibitory neurons play a determinative role in defining the population spiking state. We further examined the relationship between spiking states and classic signatures of cortical state in S1 of awake mice, finding that spiking states are highly informative of whisking activity as well as predictive of fluctuations in sensory responses. Our results demonstrate how unsupervised statistical modeling of populations of individual neuron spiking activity dynamics extends classical metrics of brain state.

## Results

### Latent state model of spiking activity

We developed a data-driven modeling framework to relate latent states of spontaneous spiking activity to LFP and whisking behavior characteristics and trial-by-trial variations in sensory-evoked responses. Recordings of the LFP and single-neuron spiking activity were acquired using linear multi-electrode arrays (Neuronexus A1×32-5mm-25-177-A32) in the primary somatosensory cortex (S1) of the awake, head-fixed mouse while monitoring whisking activity with videography (see Methods). We analyzed spiking activity from an average of 26 neurons in each recording (range 9 to 32, N= 6 recordings, see Table 1; see Methods for spike sorting metrics) distributed across layers 2 through 5 and categorized as FS (fast-spiking) inhibitory neurons and RS (regular-spiking) putative excitatory neurons as a function of their spike waveform.

**Table 1.** Log likelihood calculations under Poisson-HMM. Comparison of data (or simulated data) log-likelihoods under the Poisson-HMM, for the recorded data (1-state), recorded data (7-state), simulated data (7-state), shuffled simulated data (7-state). Simulation length is matched to recorded data length, so comparisons can be made within each row, but not directly across rows.

| Recording | Data (1-state model) | Data (7-state model) | Simulation (7-state) | Shuffled (7-state) |
| --- | --- | --- | --- | --- |
| 1-1 | -428549 | -399075 | -401893 | -437928 |
| 1-2 | -295463 | -279170 | -269299 | -287421 |
| 2-1 | -426533 | -400726 | -401244 | -431003 |
| 3-1 | -255510 | -220216 | -217860 | -252243 |
| 3-2 | -508583 | -469814 | -448926 | -500754 |
| 4-1 | -198327 | -185925 | -184549 | -200262 |

Spontaneous spiking activity in a population of FS (blue) and RS (red) neurons (10-second sample, Figure 1A) was used to fit a Poisson-emission hidden Markov model (HMM, see Methods), which has previously been used to capture coordinated firing rate changes across populations of neurons (31–34). The fitted parameters, estimated on half of the recording (Methods), are comprised of firing rates *λ_n_*(*S*^spk^) (spikes per 40-ms time bin) for each state *S*^spk^and single neuron *n* (Figure 1C) and the transition probabilities *A_ij_* between states (Figure 1D). In Figure 1, the states are ordered by the average firing rate across cells, and the cells are grouped with the FS neurons (blue) and RS neurons (red) together.

**Figure 1.**
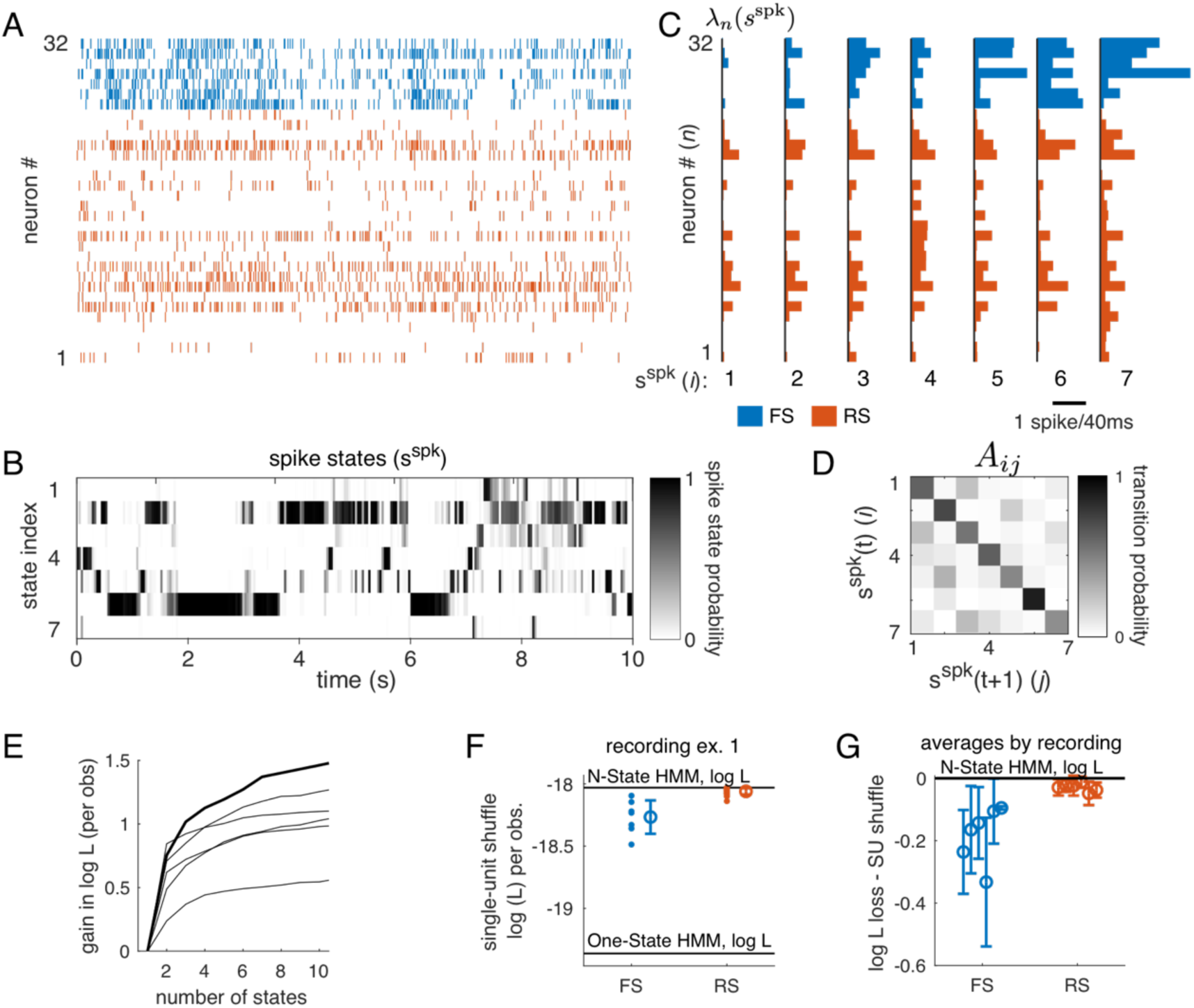
Multi-state dynamics of population spiking activity are primarily driven by FS cells. A: Spontaneous spiking activity simultaneously recorded across 32 cells in awake mouse S1. Spikes from FS cells in blue; from RS cells in red. B: States inferred by a Hidden Markov Model (HMM) during the spontaneous segment in (A). Each row is a state, and each column is a time bin. The shading (white to black) indicates the inferred probability of being in each state at each point in time. C and D: Parameters of the HMM underlying inference in panel (B). C: Firing rate parameters (*λ_n_*(*S*^spk^)) across the population of cells, including FS cells (blue) and RS cells (red). Bar length is the rate in spikes per 40-ms bin. D: Matrix of state transition probabilities (*A_ij_*). States in B, C, and D have been ordered based on average state-dependent firing rate across all cells. E: Gain in the log-likelihood relative to the one-state model for each recording (bold, example in A-D). All recordings were better fit by a model with multiple states than by the one-state (constant rate) model. Error bars are suppressed for clarity and are approximately 0.15 based on the variance across segments of the data (Methods). F: Impact of individual cells on the fit to the population was assess by shuffling spike counts across time for each single neuron, then recomputing the likelihood using the single-neuron shuffle. Single-neuron shuffle log likelihood is plotted separately for FS neurons (blue) and RS neurons (red). Top horizontal line: full-population (no shuffle) likelihood for the 7-state model (shown, A-D). G: Average single-neuron shuffle loss in the log-likelihood for all recordings. Shuffling an FS neuron decreased the log-likelihood more than shuffling an RS neuron.

Under the fitted model, we computed the probability that the population is in a particular state at each point in time, shown over the 10 second epoch for this example (Figure 1B). To get a qualitative sense of the structure captured by the model, note the “state 6” segments in the population raster shown in Figure 1A: these periods are marked by higher rates in the FS neurons (cells 26 to 32) and a few of the RS neurons (Figure 1C). Some spike-states have overall lower firing rates than others (e.g., state 1 with 0.13 spikes/bin versus state 7 with 0.54 spikes/bin), but the distinction between states also derives from differences in the pattern of firing, as quantified by the Bhattacharyya coefficient, a generalized measure of differences between multinomial distributions, which ranges from 0 (no overlap) to 1 (full overlap). For instance, state 4 and state 5 have similar average firing rates (0.29 spikes/bin and 0.36 spikes/bin, respectively), but are highly distinguishable based on spiking pattern (Bhattacharyya coefficient of 2.8 · 10^-13^; see Methods).

To assess model fit quality, we computed the likelihood on the reserved test set for models with one to fifteen states. For all recordings, a multi-state model had a higher likelihood than either the 1-state model or the shuffled model (Figure 1E, Table 1). However, it was not possible to identify a clear best choice for the number of states based on a cross-validated likelihood estimate. Thus, in subsequent analyses, we continue to show examples from the seven-state model, as after this point, the gain from additional states is minimal. All subsequent analyses were carried out for models fit with 2 to 10 states, with results shown in summary for different numbers of states.

We next examined the contribution of individual neurons to the model fit using a shuffle test. For each neuron, we shuffled the spike counts across time, thus creating a surrogate dataset preserving structure in all neurons except the shuffled cell. We recomputed the data likelihood using this surrogate set and compare the single-neuron shuffle likelihood to the original likelihood computed in Figure 1E. In the example recording (Figure 1F), we find that likelihood dropped more when an FS neuron was shuffled than when an RS neuron was shuffled, and this trend held across all recordings (p < 0.01, z = 7.0, hierarchical bootstrap over N=6 recordings, with 10^4^ shuffles, Figure 1G). This effect was not entirely explained by the higher firing rates of FS neurons relative to RS neurons, as we observed the same difference between FS and RS neurons when comparing the coefficient of variation of state-dependent firing rates extracted from model parameters (*λ_n_*(*S*^spk^); p_boot_<0.01; z = 3.1, hierarchical bootstrap). We conclude that FS neurons played a larger role within the model than RS neurons.

Finally, to assess overall model fit quality, we compared the likelihood computed for the test set to an idealized case, in which observations were generated from the fitted model. Using models (*A_ij_*, *λ_n_*(*S*^spk^), with 1 to 15 states) that were fit to the data as previously described, we simulated observations, matched to the data length of the original recording. Qualitatively, we found that a seven-state model (Figure 2A; parameters from Figure 1) exhibits coordinated fluctuations in activity observed across multiple single neurons. Such co-fluctuations and temporal modulation are missing in the one-state model (Figure 2B). In the simulation, the sequence of states (simulation ground truth, Figure 2C, top) was known, and was accurately inferred from the simulated observations (inferred states, Figure 2C, bottom). Thus, qualitatively, the multi-state Poisson-HMM captures the structure across single neurons and across time observed in the recorded data.

**Figure 2.**
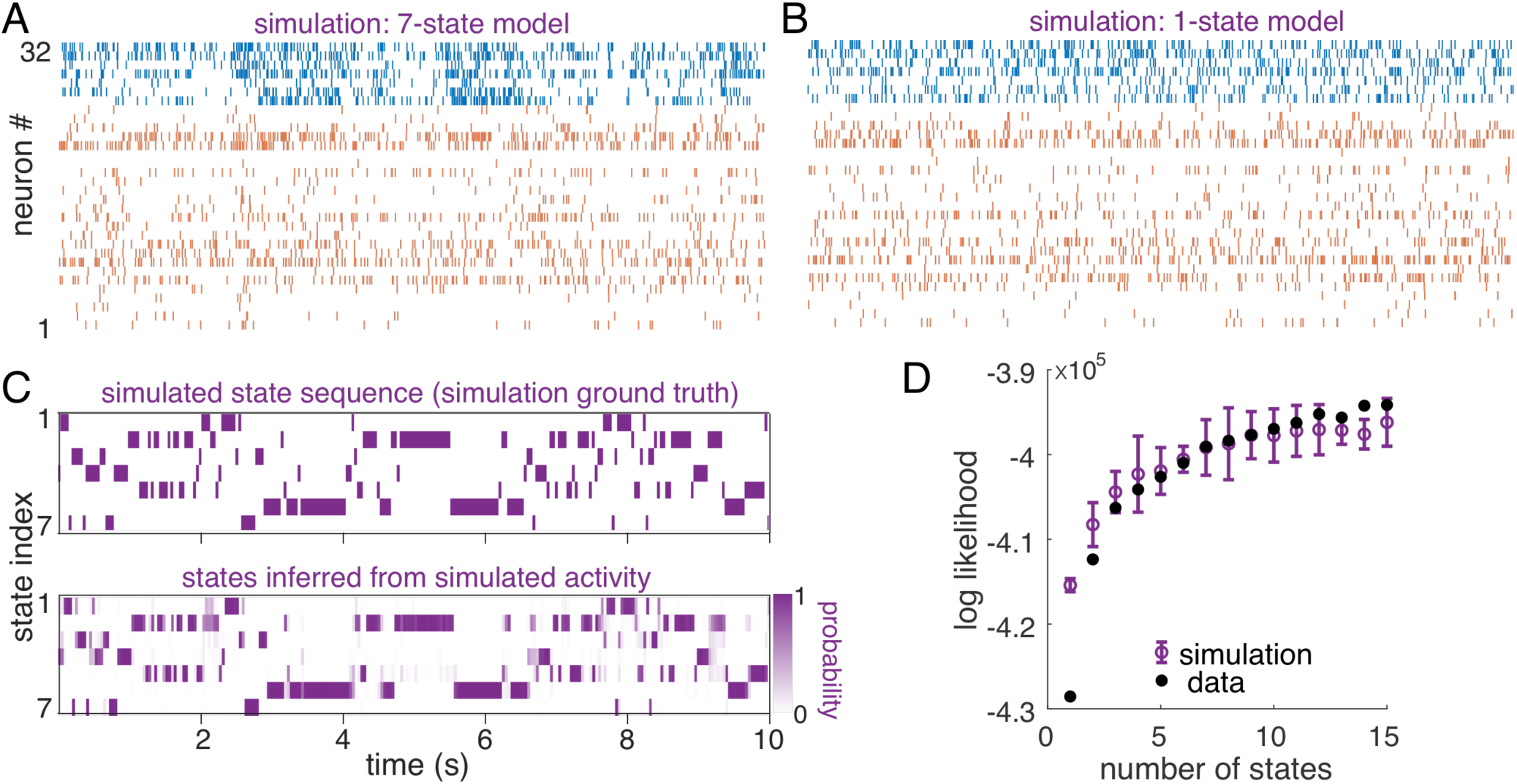
Neural data was fit by the HMM as well as surrogate data generated directly from the model. A: Seven-state model simulation for 32 cells and 10 s duration using model fit (parameters shown in Figures 1C and 1D). A sequence of states was generated from the transition matrix (Figure 1D) and observations drawn according to the emission distributions (Poisson spiking, rates in Figure 1C). B: One-state model simulation for 32 cells and 10 s duration. C: Top: the ground-truth state sequence used to generate spiking pattern (B). Bottom: inferred state probabilities, where dark purple indicates high probability. D: Log-likelihood of test-set (reserved) data under models fit with 1 to 15 states (filled black circles). Log-likelihood of simulated data, using the fitted model and matching data length (open purple symbols; error bars are the standard deviation across repetitions of the simulation).

To quantitatively assess model fits, we compared likelihood of decoded states in the data to that from ideal simulations and null tests generated by shuffling data. We first compared the likelihood of observations from simulated spontaneous S1 spiking (from the known model) to that of the recorded data (using a model fit on the reserved training set). Simulation length was matched to that of the recorded data (see Methods). We plot the log-likelihood (normalized by the number of time bins times the number of neurons) for data (filled black circles) and simulation (open purple circles) as a function of the number of states in the model (Figure 2D). For this recording, the log-likelihood of the data matched that of the simulated data for models with five states and more, indicating a fit that is as good as if the model were the “true” generating model for the data.

### Relationship of LFP and spiking states

We next quantified the relationship between spiking states and the LFP, using an established metric for defining ‘states’ of the LFP (22). Here LFP state is determined by calculating a power spectrogram of the LFP (Methods) and comparing the power in low-frequency (LF, 1-10 Hz) components to the power in high-frequency (HF, 30-90 Hz) components (Fig. 3B). We refer to these as “LFP state L” (LF-dominant, with high LF/HF, green, Figure 3A, B) and “LFP state H” (HF-dominant, with low LF/HF, black, Figure 3A, B). The spiking states inferred during a segment of LFP state H (Figure 3C) and LFP state L (Figure 3D) are shown with the population spike raster overlaid. Transparent overlay (graded from black to green) represents distinct spiking states during the recording intervals for which spike-state was assigned with >80% probability. This example illustrates a tendency toward observing spiking states 1 and 2 during LFP state H (Figure 3C) and a tendency toward observing spiking states 5, 6, and 7 during LFP state L (Figure 3D).

**Figure 3.**
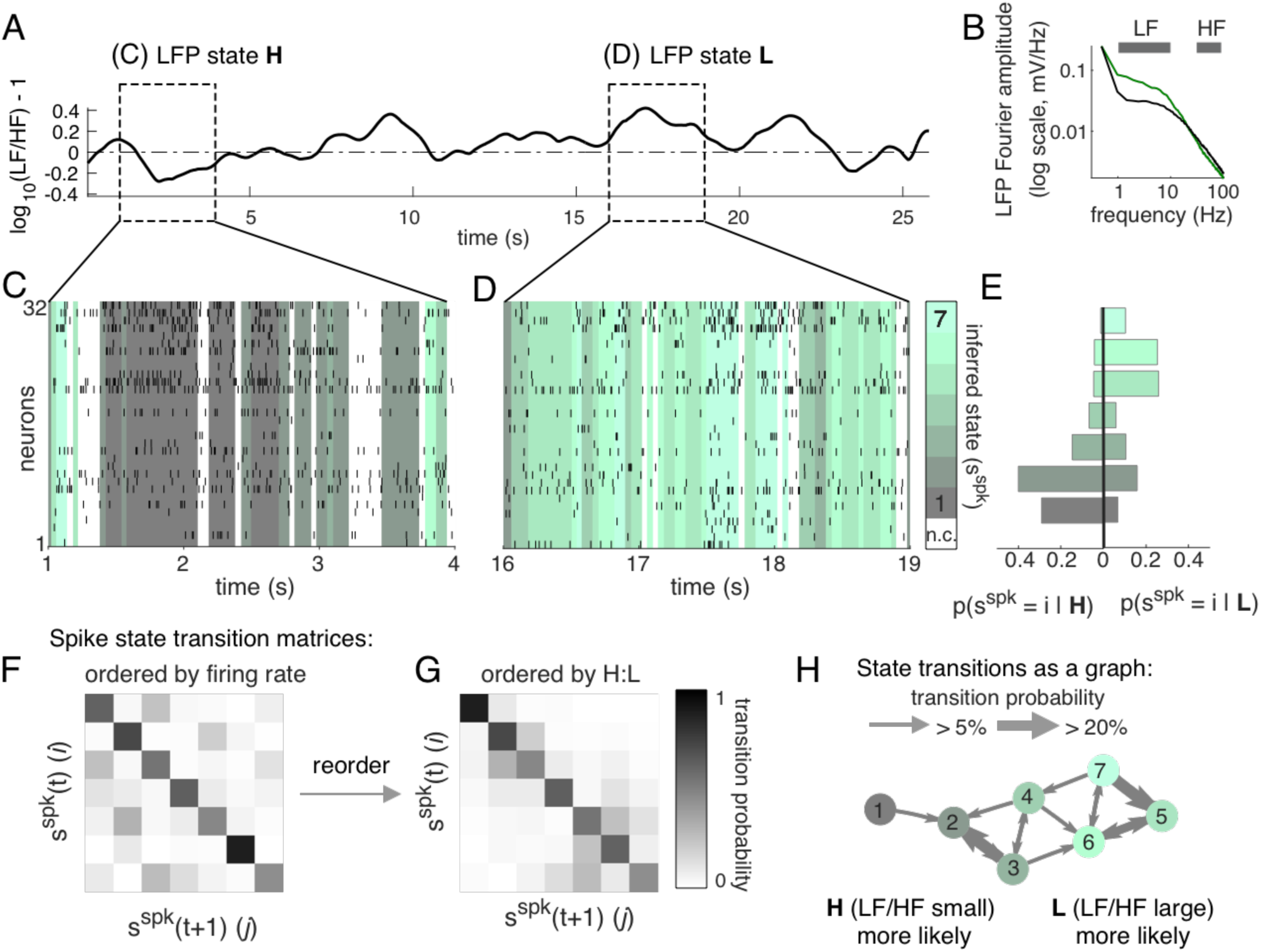
LFP states are coordinated with distinct states of spiking activity. A: Example period of spontaneous activity in which LFP state transitioned from state H to state L, defined using Fourier amplitudes spectrum features (B). B: Fourier spectrum of the LFP, showing the LF frequency band (1-10 Hz) and HF frequency band (30-90 Hz). Division into states (L, green; H, black) is by comparing average amplitude in LF to that in HF (“LF/HF”), following previous studies (22). C - D: Inferred spiking states (colors) with overlaid spike raster during segment 1 (LFP state H, panel C) and segment 2 (LFP state L, panel D). White indicates state probability was less than 80%. E: Distribution of spiking states observed during LFP state H (left) and during LFP state L (right). Spiking states 1 and 2 are more likely during LFP state H, while spiking states 5, 6, and 7 are more likely during LFP state L. F: The transition matrix (*A_ij_*, reproduced from Fig. 1D) ordered by firing rate. G: Transition matrix re-ordered using relative fraction of time the LFP state was H or L while spiking activity was inferred to be in state *i.* Reordered transition matrix (right) has nearly block-diagonal structure. H: Directed graph illustrating transition matrix. Common transitions (*A_ij_* > 0.05) between states (circles, color as in B and C) are indicated by directed arrows between nodes (states) in the graph. Self-connections (*A_ii_*) not shown.

To quantify the tendency of specific spiking states to occur more often during LFP state H or L, we divided the recording based on LFP state, then computed the histogram of inferred spiking state restricted to state H (Fig. 3E, left) or state L (Fig. 3E, right). The color map used for spiking states in Fig. 3 was ordered based on the LFP, using the ratio of time each spiking state was in LFP state H to LFP state L. This ordering differs from that used in Figure 1, in which states were sorted by average firing rate, which produced a transition matrix with many off-diagonal elements (Figure 3F). In contrast, LFP-based sorting revealed nearly block-diagonal structure in the spiking state transition matrix (Figure 3G). We visualized the spiking state transition matrix as a directed graph (Figure 3H), with a node for each state and in which the thickness of the arrows between spiking states represents the probability of a transition. Spiking state 1 is directly connected to spiking state 2 (both more likely in LFP state H) but moving from spiking states 1 and 2 to states 5, 6, and 7(more likely in LFP state L) requires multiple steps. This indicates that while traversing the graph according to the transition matrix probabilities, the system will tend to stay in LFP-H spiking states or LFP-L spiking states for long periods of time.

Based on the tendency for spiking states to align with one LFP state or the other, we expected that using the state representation of spiking activity would predict the LFP state (H or L). Thus, we built a binary classifier (Figure 4) to predict the LFP state (H or L) based on either direct spiking metrics (single neuron spike count, summed population spike count, or population vector) or on the inferred spiking states obtained using the HMM. Because LF/HF required a 1-second window to compute, for this analysis 40-ms time bins were sampled at 1 Hz across the experiment, then divided randomly into training and test sets, balancing the number of observations associated with each LFP state so that chance-level performance was 50%. For decoding, we used an SVM classifier with a linear kernel (see Methods).

**Figure 4.**
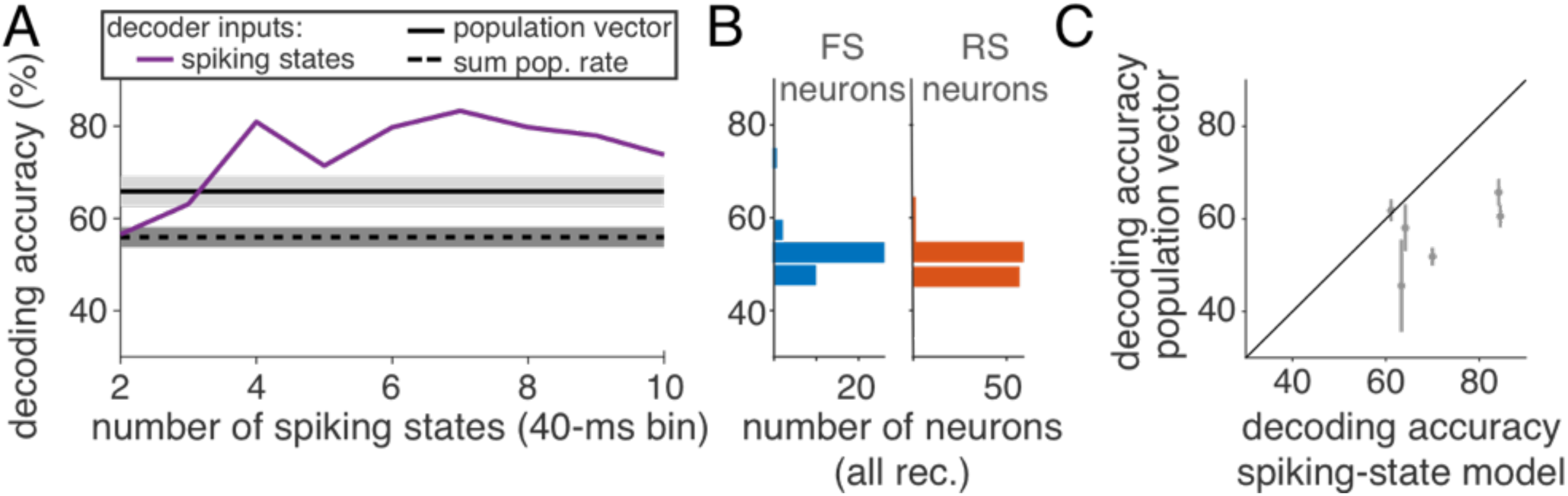
Decoding LFP state from spiking activity. **A.** Decoding accuracy (chance level, 50%) of determining LFP state (H or L) from inferred spiking states (purple), population vector (solid black), and summed population rate (dashed) for an example recording (1–1). **B.** Decoding accuracy, using single neuron activity (FS, left/blue; RS right/red), across all neurons. **C.** Across 5 of 6 recordings, decoding accuracy is higher from inferred spiking states than from the population vector.

To examine what aspects of population activity were informative of LFP state, we built a series of decoders. In the simplest decoder, the predictor was the summed spike counts across the full population (single recording example, “summed population rate”, Figure 4A, dashed black line), which was a poor predictor of LFP state (accuracy, 54% ± 5%, range across 6 recordings 46% to 59%). To utilize the structure in the population firing pattern, we next constructed a linear classifier based on the population rate vector. The population vector classifier performed better than the summed-activity classifier (Figure 4A, “population vector”, solid black line), with performance in this example recording of 66% ± 3% (across cross-validation folds). Finally, we predicted LFP state using as input the spiking state inferred under models having from 2 to 10 spiking states (Figure 4A, “spiking states”, solid purple line). The highest decoding performance was achieved by models with at least 4 states, above which decoding performance leveled off.

To determine whether the accuracy of decoding LFP state was explained by a minority of high-performing cells, we used as predictors spike counts from single FS or RS neurons (Figure 4B). While a few FS neurons were informative of LFP state (accuracy of best FS in each recording, 58 ± 7%, range 53% to 70%), most neurons decoded LFP state poorly (average decoding accuracy: 50% ± 2% (RS), N = 119; 52% ± 4% (FS), N = 39). Across five of six recordings, the best-performing decoders used inferred spiking states (Figure 4C), achieving an average accuracy of 71% ± 10% (n = 6 recordings, range 61% to 85%). Figure 4C shows the decoding accuracy using 7 spiking states. In comparison, decoding with the population vector was lower by 14 percentage points (n = 6 recordings, 57% ± 7%).

### Spiking states are predictive of trial-by-trial variability in sensory-evoked responses

Given previous reports of the predictive nature of ongoing LFP activity on sensory evoked responses in this pathway (16), the relationship between spiking states and the LFP suggests similar predictive capabilities through the latent spiking states. We now directly examine the extent to which latent spiking states are predictive of trial-by-trial variability in sensory-evoked responses in single neurons.

Between periods of spontaneous activity, we recorded sensory-evoked responses to simple punctate single-whisker stimuli as described previously (16,35). We first established that the state of pre-stimulus activity, classified under the spiking state model developed for spontaneous activity (Figure 1), was predictive of single-trial evoked responses. We quantified the sensory-evoked responses, conditioned on the pre-stimulus state, and identified neurons with state-dependent sensory-evoked responses at a statistically significant level (see Methods). Briefly, using one second (1 s) of pre-stimulus activity, we classified each trial based on the pre-stimulus spiking state that was the most probable state immediately prior to stimulus delivery. Trials were partitioned based on pre-stimulus spiking states, with population rate parameters plotted in Figure 5A and sensory responses plotted in Figure 5B. For comparison, black-outlined bars show the peri-stimulus time histogram (PSTH) averaged across all trials (Figure 5B). Qualitatively, in some states (state 1, Figure 5A, B) the sensory response was less than in other states (e.g., state 4, 5), both in terms of the absolute number of spikes in the post-stimulus window and the size of the evoked (i.e., subtracting the pre-stimulus spike count) response (Figure 5B).

**Figure 5.**
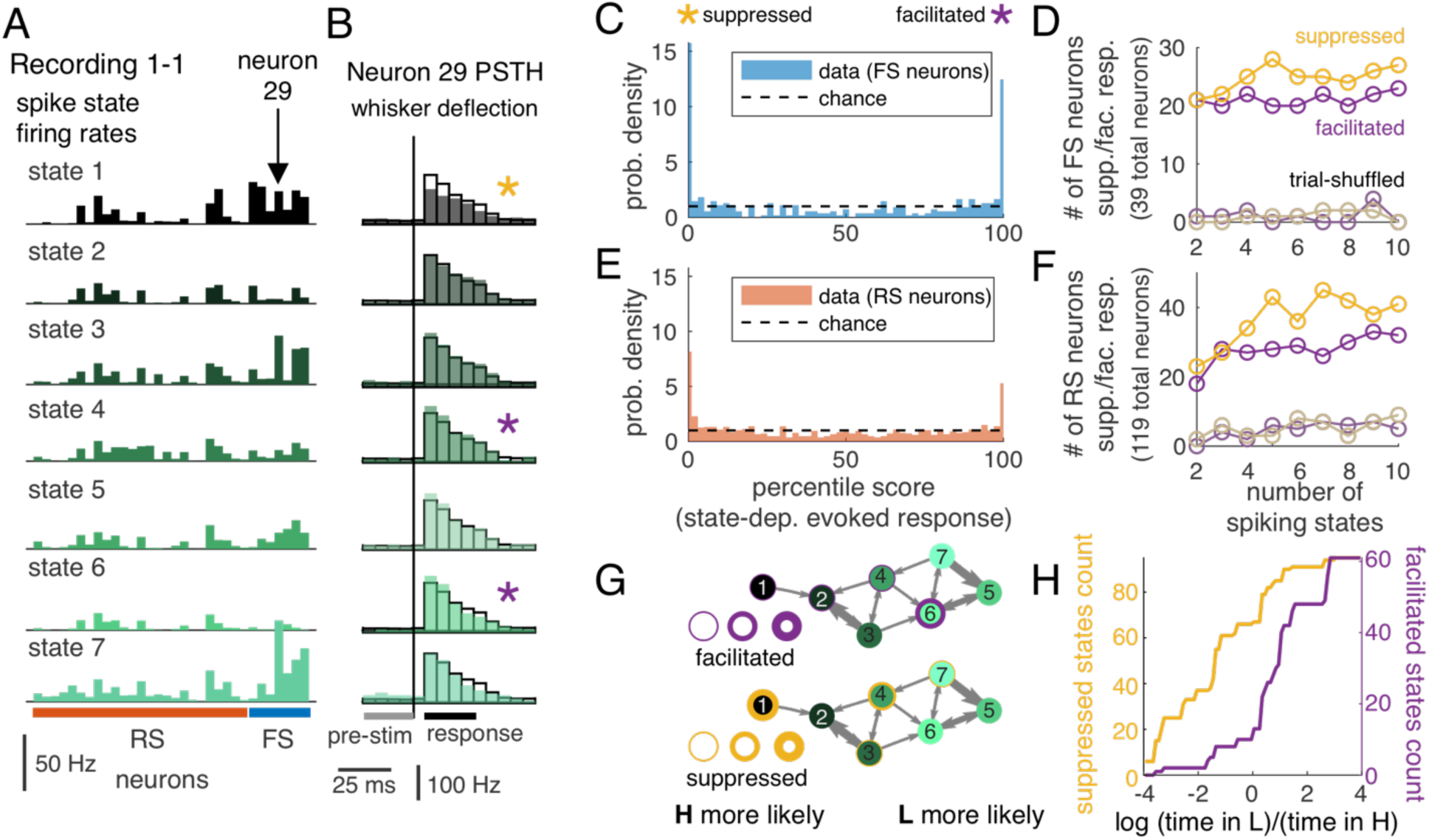
Population spiking states are predictive of variability in sensory responses in single neurons. A: Patterns of firing rates across neurons in each state. Bars represent the model rate parameter (spikes per second) in single neurons in a 40-ms bin (as in Figure 1). State colors as in Figure 3. B: Responses to whisker stimulation (punctate deflection at time indicated by gray line) when trials are sorted by the decoded pre-stimulus state. The single neuron shown as an example in B is marked in panel A. PSTH bin width is 5 ms. Significant state-dependence is indicated by a purple star (facilitated) or yellow star (suppressed). This is defined relative to the sensory response averaged across all trials (black outline) based on p < 0.01 in a 2-sided shuffle test (Methods). C: Distribution of percentile values for state-averaged responses for FS neurons from all recordings for pre-stimulus-sorted (color). Black dashed line shows uniform (chance-level) distribution of percentile values. D: Number of FS neurons with detected state-dependent responses using models with increasing numbers of states. Yellow/purple indicate suppressed/facilitated. Trial-shuffle control is shown in light colors. E: Same as C, for RS neurons from all recordings. F: Same as D, for RS neurons from all recordings. G: State transition graph (as in Figure 3), with purple/yellow indicating presence of facilitated / suppressed evoked responses in that state. Thicker outline represents more neurons with significant state dependence. Table 2 reports counts of facilitated/suppressed neurons for each recording. H: Cumulative count of observed values of ratio of time spent in LFP state L to time spent in LFP state H for spiking states identified as suppressed (yellow) and facilitated (purple). Note the log scale, so negative (positive) values are states more frequently associated with LFP state H (state L). States with suppressed responses tended to be associated with LFP state H. Totals do not match the neuron counts in D and F because some individual neurons had facilitated (or suppressed) responses in more than one state.

**Table 2.** Number of state-dependent evoked responses detected in each recording, under the 7-state model.

| Recording<br>(total count) | 7-state model<br>facilitated count | 7-state model<br>suppressed count |
| --- | --- | --- |
| 1-1 (32) | 12 | 14 |
| 1-2 (32) | 9 | 12 |
| 2-1 (28) | 5 | 8 |
| 3-1 (11) | 7 | 12 |
| 3-2 (36) | 19 | 34 |
| 4-1 (19) | 9 | 15 |

To quantify significance of apparent state-dependence in sensory responses of single neurons, we used a shuffle test. A distribution of average response size by state was generated by randomly assigning trials to each state. The empirical average state-dependent sensory response was then compared to the shuffle distribution, represented as the percentile of the magnitude of the state-dependent average evoked response within the boot-strapped distribution (FS neurons, Fig. 5C, and RS neurons, Figure 5E; in a 7-state model). For reference, if there were no dependence of sensory response on the spiking state, the distribution of percentiles would be uniform (dashed black lines, Figure 5C, 5E). Based on this percentile value, state-dependent responses were categorized as “suppressed” (percentile <0.5; yellow star) or “facilitated” (percentile >99.5; purple star), or not significantly different from the distribution of responses across all trials. Across the population of neurons, more than half of neurons (for 7-state mode: 91 of 158 total, range 8 to 24 per recording, from N = 6 recordings) had at least one state-dependent sensory-evoked response, exceeding the false-positive rate expected by our criterion for rejection (1%).

The organization of state-dependence across cells was non-random, with state-dependent responses more common in the FS population than the RS population. For instance, using the 7-state model, 58% of all FS neurons were facilitated in at least one state (range across individual recordings: 37% to 80%, N = 6 recordings) and 70% of all FS neurons were suppressed in at least one state (range: 37% to 100%, N = 6 recordings), while 21% of RS neurons were facilitated (range: 9% to 31%) and 39% of RS neurons were suppressed (range: 23% to 54%). For this analysis, cells were only counted once, so if there were two states with suppressed responses (as in Figure 5B), the neuron contributed once to the count in Figure 5D. We note that, while we show the seven-state model for illustration, this choice was not critical for the detection of state-dependent responses. Models with four or more states show a relatively constant proportion of neurons with state-dependent responses (Figure 5D, F). We verified that this is not simply a consequence of limited trial numbers by repeating the analysis for trial-shuffled controls with matched trial counts per state (gray, “trial-shuffled”, Figure 5D, F).

Finally, we examined the extent to which specific pre-stimulus spiking states were associated with facilitated or suppressed responses. As in Figure 3, we show state transition graphs, annotated to show the organization of facilitatory and suppressive pre-stimulus states. Rings around each state indicate the fraction of single neurons with either facilitated (Figure 5G, purple) or suppressed (Figure 5G, yellow) responses, relative to the average evoked response across all trials, where thicker rings represent more neurons with significant state dependence. Suppressed states (yellow) were consistently found in the LFP-state-H associated spiking states (black), and a subset of LFP-state-L spiking states (lighter green) were associated with facilitated responses (purple). Across all recordings (Figure 5H, using 7 spiking states), facilitated spiking states were more likely to be observed during LFP state L (median time ratio, 1.6), while suppressed spiking states were more likely to be observed during LFP state H (median time ratio, 0.6).

### Spiking states track whisking activity

During wakefulness, LFP state (quantified by LF/HF) is correlated with whisking behavior (5). Thus, we expect that inferred spiking states are informative of whisking behavior as well, illustrated by aligned trajectories of LFP (Figure 6A), spiking states (Figure 6B), and whisking behavior (Figure 6C) over an example 25-second period. As in Fig. 3 and 5, spiking states are ordered based on the LFP state observed but utilizing a color map to enable easier discrimination between spiking states. Spiking states 1 and 2 are those for which low LF/HF is most likely. In the example shown in Fig. 6B-C, spiking states 1 and 2 (dark blue shades) appear during bouts of whisking (Fig. 6C), but rarely otherwise, suggesting that spiking state dynamics were closely tied to whisking behavior. Across the full recording, the onset of whisking coincided with an abrupt transition of the inferred spiking states into spiking states 1 or 2 (post-onset state is 1 or 2 in 25 of 38 detected onsets; Fig. 6D). By comparison, we observed a much slower decrease in LF/HF (Fig. 6E).

**Figure 6.**
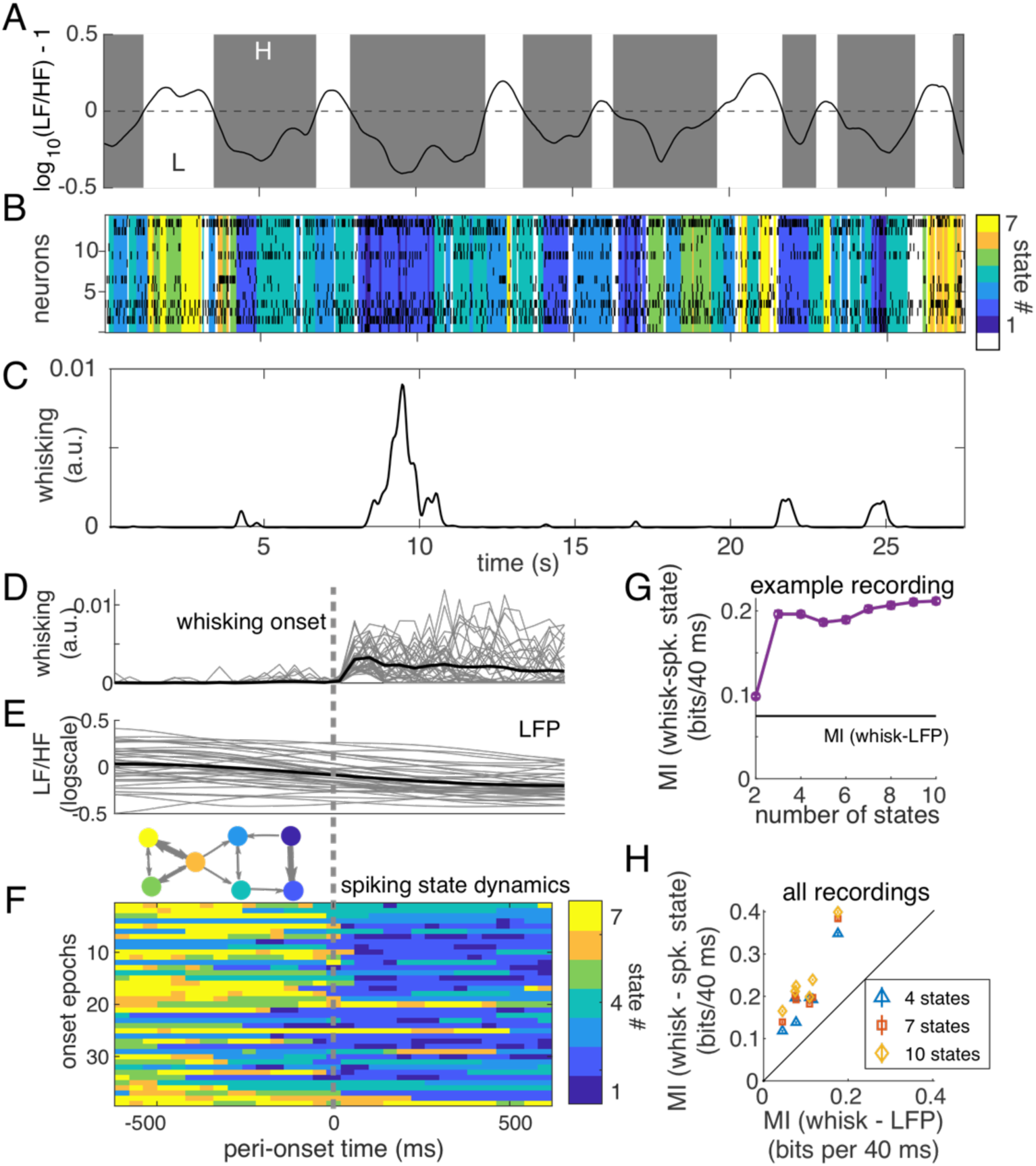
Spiking state dynamics are related to whisking activity. A: LFP state metric (log_10_ (LF/HF) – 1) during spontaneous activity. Gray shaded regions show where LF/HF < 10 and white for LF/HF > 10. B: Spiking activity across 15 neurons and the inferred latent state during the same period of spontaneous activity. Colormap (differing from Figure 3, 5) was chosen for higher contrast between states. C: Whisking activity, quantified from motion energy (frame-to-frame changes, see Methods) in videography. D-F: Onset of whisking is accompanied by a gradual decrease in LF/HF and rapid change of spiking states. D: Onset times were identified using whisking energy, requiring a period of no whisking over 400 ms followed by crossing a threshold. E: LFP state metric (log_10_ (LF/HF) – 1) gradually decreases at the onset of whisking. Gray lines: individual events. Solid black is the average. F: Spiking state dynamics before and after onset of whisking, for 38 detected whisking events. State transition graph is shown for reference. G: For this example, information between spiking state representation and whisking activity, for state models with 2 to 10 states. Information carried by comparing LF/HF to 10 is marked by the black line. H: Information about whisking activity calculated on all six recordings for 4-state, 7-state, and 10-state models are all higher than the information carried by LF/HF > 10. Error bars estimated by resampling half of the data and repeating information calculation (50 repeats) and are typically smaller than the markers.

To quantify this observation across datasets, we calculated mutual information (MI) between whisking activity and (a) LFP states (based on LF/HF) and (b) the inferred spiking states (see Methods). For the example recording in Figure 6D, we found that the spiking states inferred using a model with at least three states were more informative of whisking/non-whisking behavior (0.25 ± 0.02 bits/bin, whisking-spike states, bin of 40 ms) than the LFP state was (0.15 ± 0.02 bits/bin, Figure 6G). In all recordings, inferred spiking states (with at least 4 states, showing 4, 7, and 10 states in Figure 6E, “whisk - spiking states”) were more informative of whisking activity than the LFP state (“whisk – LFP”), as shown in Figure 6H. We conclude that modeling the latent dynamics of spontaneous population spiking activity provides a useful metric of cortical state in awake mouse S1 that is related to traditional LFP-based metrics and contains information about the representation of sensory information and changes in behavior with high temporal resolution.

## Discussion

Here we applied a latent state modeling framework to recordings from the awake mouse somatosensory cortex and compared the latent states inferred from population spiking activity to standard metrics of brain state. We found that latent spiking states captured structure in population spiking activity and were closely linked to slow-changing LFP states, predictive of fluctuations in single-trial sensory-evoked responses, and informative of whisking activity. Fast-spiking neurons had higher impact on the latent state than regular-spiking neurons, reflecting a large role of inhibitory cell activity in determining state.

### Latent dynamics of population spiking activity in mouse S1

The latent-state representation of population activity inferred using a Poisson-HMM was highly informative of the state of wakefulness, quantified by the standard measures of LF/HF and whisking behavior. Spiking states were even more informative of whisking activity than the LF/HF metric. Moreover, LFP states (based on LF/HF) were more accurately decoded from the latent state representation of spiking activity than they were from more direct measures, such as the summed firing rate across the population or even the population rate vector. The improvement of the inferred spiking states from the HMM over the population rate vector likely arises due to the temporal information embedded in the inferred spiking state, indicating the importance of accounting for correlations in time for analysis of spiking data.

We also found that inferred spiking states were predictive of single-trial sensory-evoked responses. In past work, we built a framework for predicting single-trial sensory-evoked LFP responses in awake mouse S1, in which features of the pre-stimulus LFP recording were identified based on their ability to predict response characteristics (16). There, we found that our most predictive LFP features were equivalent to LF/HF, which had been used previously as a metric of state in S1 (5,22). Thus, in the current study, we expected that pre-stimulus spiking states would also be predictive of single-trial sensory-evoked responses in single neurons, and that is what we found (Figure 5). The tendency of the “suppressed response” pre-stimulus states to align with low LF/HF is consistent with the predictive features examined in our previous study, which found smaller LFP responses in a low-LF/HF state. Thus, the Poisson-HMM framework produces a representation of population dynamics that captures enough of the spatial (across cells) and temporal (across time) structure in cortical activity to enable accurate decoding of the LFP state and whisking behavior and to predict single-trial fluctuations in sensory responses in single neurons.

### FS neurons and population spiking state in mouse S1

We found that activity of FS neurons, which are more likely to be inhibitory cells (36), was more strongly linked to the overall population spiking state than the activity of RS neurons, beyond any dependence based on firing rate alone. For instance, model fit quality degraded substantially more when the temporal structure of FS neuron activity was disrupted compared to that of RS neuron activity. Further, a larger fraction of FS neurons than RS neurons exhibited state-dependent sensory response patterns. Our general finding that spiking activity in inhibitory neurons is tightly linked to population spiking state and, by extension, the LFP, is consistent with past work focused on single-neuron contributions to the LFP that suggested that LFP primarily reflects inhibitory activity (37). There is variability across single neurons within FS and RS classifications, however: for example, not all FS neurons are tightly linked to the latent state. Understanding the circuit dynamics that underlie the emergence of population spiking states will require finer description of the cell-type specific roles, possibly enabled by new datasets with more specific cell-type identity information (38) and by computational models permitting cell-type classification based on single-neuron spiking statistics (39).

### Insights from a simple latent-state model

In the Poisson-HMM used here to model spiking states, all cells were assumed to be linked to a single hidden state at any moment of time, and this state is assumed to have Markovian dynamics, with dependence only on the current state. While these assumptions limit the flexibility of the model, inferred spiking states from the model were successful in predicting LFP state, sensory-evoked responses, and whisking activity. Several previous studies have used the Poisson-HMM framework to describe population activity patterns and dynamics (10,33,40,41). For instance, the Poisson-HMM captured the structure of fast sequences of population activity patterns in rat gustatory cortex, demonstrating step-like rather than ramping behavior during a sensory-decision-making task (40). We used a coarser time bin, lasting 40 ms rather than 1 – 2 ms in the Jones study. We chose this bin size because our analysis concerned spontaneous activity over relatively long (minutes) segments of data. Thus, our aim in using the Poisson-HMM is to obtain a relatively simple representation of spatial and temporal dynamics. The same model was also used to capture spontaneous activity states in the five seconds preceding a sensory-evoked response (33), and there it was argued that the identified states reflect underlying attractor dynamics. Due to the prevalence of FS neurons in the spiking patterns that determined spiking states, a different interpretation of states may be more appropriate in our study. An intriguing idea, raised by a recent modeling study, is that PV interneurons are poised to switch the cortical network in and out of an inhibition-stabilized network (ISN) dynamical state (42). Such state switches would be reflected in differences in sensory-evoked responses to well-controlled stimuli, as we observed, suggesting that changes in ISN dynamics may parsimoniously account for the state-dependent patterns of sensory-evoked responses we observed.

### Limitations of Poisson-HMM for spiking dynamics

While the Poisson-HMM has the advantage of a small number of parameters and simple state dynamics, the assumption that all neurons are in the same state likely breaks down in populations spanning larger regions of cortex. An alternative approach, more commonly used in non-sensory brain areas (motor, prefrontal, hippocampus) is to embed neuronal activity patterns in a low-dimensional manifold upon which population activity smoothly evolves. For example, latent structure in task-driven (non-spontaneous) population dynamics inferred utilizing a dynamical system improved decoding of movement intention (43), and other studies have identified low-dimensional structure in prefrontal cortex (44) and hippocampus (45). Responses in visual cortex to visual inputs appear to live in a distinct subspace from spontaneous activity (28). Determining which latent state model to use in which recording would be clarified by developing rigorous “null” computational models, in which latent dynamics and population responses can be carefully controlled (46,47).

### Future directions

Understanding of the dynamics of local brain state has advanced tremendously through careful dissection of modulatory circuit mechanisms in primary sensory areas. Experiments utilizing patch-clamp electrophysiology in awake mice have revealed state-dependent changes in the activity of single neurons of specific cell classes (15,25), shared patterns of variability across small numbers of cells (48), and control of excitability through specific cell type activation (49). In the large-scale multi-single-neuron recordings enabled by modern high-density multielectrode recordings, such fine-scale anatomical information is rarely available. On the other hand, highly parallel recording methods reveal functional relationships across the population that are not observable from recordings of single neurons, and may carry rich information on the functional roles of specific anatomical cell types. The latent dynamics uncovered by these approaches may also provide functionally meaningful and pragmatic targets for control of circuit and network activity as tools for applying control theory to neural circuits continue to emerge (50,51). A clearer understanding, through quantitative frameworks such as that presented here, of these features of cortical dynamics will ultimately provide an invaluable tool to map individual-specific differences in the neural signatures of behavior, perception, and cognition.

## Materials and Methods

### Experimental data collection

Data presented here were previously reported in (35) and (16) and is available. All procedures were approved by the Institutional Animal Care and Use Committee at the Georgia Institute of Technology and agreed with guidelines established by the National Institutes of Health. Four nine-to twenty-six-week-old male C57BL/6J mice were used in this study. Mice were maintained under 1-2% isoflurane anesthesia while being implanted with a custom-made head-holder and a recording chamber. The location of the barrel column targeted for recording was functionally identified through intrinsic signal optical imaging (ISOI) under 1-1.25% isoflurane anesthesia. Recordings were targeted to B1, B2, C1, C2, and D2 barrel columns. Mice were habituated to head fixation, paw restraint and whisker stimulation for 3-6 days before proceeding to electrophysiological recordings.

#### Electrophysiology

Local field potential and spiking activity were recorded using silicon probes (A1×32-5mm-25-177, NeuroNexus, USA) with 32 recording sites along a single shank covering 775 μm in depth. The probe was coated with DiI (Invitrogen, USA) for post hoc identification of the recording site. The probe contacts were coated with a poly(3,4-160 ethylenedioxythiophene) (PEDOT) polymer (52) to increase signal-to-noise ratio. Contact impedance measured between 0.3 MOhm and 0.5 MOhm. The probe was inserted with a 35° angle relative to the vertical, until a depth of about 1000 μm. Continuous signals were acquired using a Cerebus acquisition system (Blackrock Microsystems, USA). Signals were amplified, filtered between 0.3 Hz and 7.5 kHz and digitized at 30 kHz.

#### Recording of spontaneous and sensory-evoked activity

Mechanical stimulation was delivered to a single contralateral whisker corresponding to the barrel column identified through ISOI using a galvo motor (Cambridge Technologies, USA). The galvo motor was controlled with millisecond precision using a custom software written in Matlab and Simulink Real-Time (Mathworks, USA). The whisker stimulus followed a sawtooth waveform (16 ms duration) of various velocities (1000 deg/s, 500 deg/s, 250 deg/s, 100 deg/s) delivered in the caudo-rostral direction (53). Whisker stimuli of different velocities were randomly presented in blocks of 21 stimuli, with a pseudo-random inter-stimulus interval of 2 to 3 seconds and an inter-block interval of a minimum of 20 seconds. The total number of trials across all velocities presented during a recording session ranged from 196 to 616. Recordings were acquired in 10-minute segments, and the end of each 10-minute segment was 1 minute of spontaneous activity. All spontaneous segments of 10 seconds or longer were analyzed.

#### LFP pre-processing

For analysis, the LFP was down-sampled to 2 kHz. Using the same quality criterion as previously (16), we screened for channels that drifted or had excessive 60-Hz line noise. Only layer-4 LFP was analyzed in this study. Layer 4 was determined using the sensory-evoked MUA activity and current source density profiles as described previously (16,35).

#### Spike sorting and single unit quality criteria

Spike sorting was performed using Kilosort2 (54) with manual curation of spike clusters in Phy (55). The quality criteria for identifying single unit activity were the same as in (35), summarized by these six requirements: (i) units had more than 500 spikes, (ii) spike waveform SNR was greater than 5, (iii) the trough-to-peak amplitude of the unit had a coefficient of variation less than 0.2 averaging over 120-s windows, (iv) firing rate (over 120-s windows) had a coefficient of variation of less than 1, (v) fewer that 1% of inter-spike intervals were shorter than 2 ms, and (vi) the Maholonobis distance between the unit waveform and the closest waveform that was *not* assigned to that unit was larger than 55 in the space of the first three principal components. Units were classified as RS or FS based on the time elapsed from the waveform trough to waveform peak, with values larger than 0.5 ms classified as RS units and those smaller than 0.4 as FS units (14,22). Intermediate waveforms (about 3% of units) were not included in the analysis. Overall, we analyzed 39 FS units and 119 RS units across six recordings (average of 26 units per recording, range 19 to 36) in four different mice. Units were distributed across layers 2 to 5, with the large majority in layer 5 (L2/3, 13; L4, 24; L5, 117; L6, 4), precluding layer-dependent analyses in this study.

#### Whisking quantification

Video recordings of whisking activity were acquired under infrared illumination at 25 Hz using a camera (6.8 pixels/mm, HQCAM) positioned below the mouse’s head, acquiring a view of whiskers as well as of the galvanometers used for stimulus control. For each movie, the portion of the field of view including the whiskers was extracted. Whisking activity was extracted by first identifying motion components using non-negative matrix factorization of the square difference between frames. Up to 20 components were identified and these were manually scored as representing whisker motion, nose motion, or galvanometer motion. Galvo traces were used to verify temporal alignment of movies and electrophysiology. Whisking activity was quantified as the sum over whisker-motion components.

#### Whisking onset analysis

To identify the transition from non-whisking to whisking behavior (analyses in Fig. 6D-F), we analyzed the whisking motion time series, obtained by summing over projections of the whisking moving into NMF modes described above. A fixed threshold for whisking was identified for each recording. To identify a transition, we required 400 ms of non-whisking followed by a minimum of two frames of whisking behavior. During the 400 ms of non-whisking behavior, at least 90% of frames (9/10 frames) had to be below the detection threshold. Transition points occurring within 500 ms of the beginning or end of a recording segment were discarded.

#### Experimental design and statistical analysis

Multiple statistical analyses were used in this study and are described in detail below. All statistical analyses were conducted in MATLAB 2019b and 2022a (Mathworks).

### Cortical State and Spiking State Model Fitting and Validation

#### Cortical state characterization

To describe cortical state in S1 of awake mice, we calculated LF/HF and separated recordings into periods with high (greater than 10) and low values of LF/HF, following a convention used in earlier studies (5,22). Specifically, we compute the spectrogram at 1-s intervals and defining LF (HF) as the average Fourier amplitude between 1 and 10 Hz (30 and 90 Hz, excluding 58 to 62 Hz).

#### Hidden Markov Model of cortical spiking activity

To describe cortical spiking states, we fit a Hidden Markov Model with Poisson emission distribution (31,33,34,40). This model assumes that the next state *S*^spk^(*t* + Δ) ∈ [1,2, … *N*_*S*_] only depends on the current state *S*^spk^(*t*), and this is characterized by the transition matrix *A*:

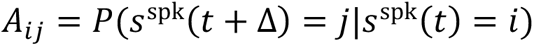

We assuming firing is Poisson and conditionally independent across neurons. Specifically, the distribution of spike counts {*y_n_*} over a time bin of duration Δ (Δ = 40ms) across the population (*n* = 1 … *N_C_* neurons) while in spiking state *S*^spk^, is given by

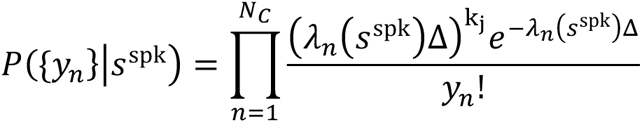

where each state *S*^spk^ ∈ {1, 2, … *N_S_*} is characterized by rate parameters *λ_n_*(*S*^spk^). The joint probability across a set of *C* simultaneously recorded cells is

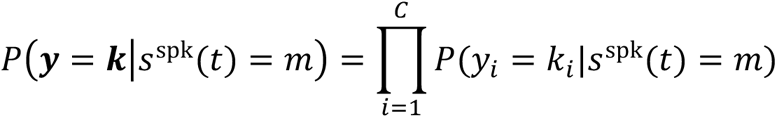

Thus, each state is defined by a vector of firing rates *λ_n_*(*S*^spk^). We binned spike counts at a temporal resolution of 40 ms to match the resolution (25 Hz) of video monitoring of whisking activity. In other works, the temporal resolution of spiking states has ranged from 1 ms (40) to 500 ms (31).

The model was implemented by adding a conditionally independent Poisson observation model to the pmtk-3 (probabilistic models toolbox, https://github.com/probml/pmtk3). To fit the model, we split each recording into two halves as follows. A recording consisted of a set of 8 to 12 segments, each of 10 minutes duration. Each segment had 10 periods of spontaneous activity (duration, 10 s to 70 s) with no controlled whisker stimulation, interspersed with periods of controlled whisker stimulation during which evoked responses were obtained. Model parameters were estimated on a training set consisting of spontaneous periods from half of the segments (1^st^, 3^rd^, 5^th^, etc.). Using the estimated model for each recording, states *S_i_*(*t*) at time *t* in each spontaneous period in the test set (the remaining half of long segments: 2^nd^, 4^th^, 6^th^, etc.) were decoded by estimating *p*(*S_i_*(*t*)|{*y*(1: *T*)) using standard methods (forward-backward algorithm). Reported likelihood values are computed across all spontaneous periods in the test set. We used two controls to interpret the data likelihood under the fitted model. First, to estimate the ceiling of model performance, we simulated an HMM using the fitted state transition matrix and drew observations from the emission distribution at each point in time according to the simulated state. We then computed the likelihood of these simulated observations under the model. Second, we verified that we could not obtain the same result in shuffled data. Thus, we shuffled the simulated observations in time and recomputed the likelihood of shuffled observations under the fitted model. Likelihood calculations for all recordings are listed in Table 1.

For the analysis of pre-stimulus state, we extracted the sensory-evoked from all segments, and using the 500-ms pre-stimulus time window, we inferred HMM state using the model previously fit on the spontaneous periods. The pre-stimulus state was assigned to the state with the highest probability in the final pre-stimulus time bin (−40 ms to 0 ms pre-stimulus).

#### Quantifying differences between spiking states

To analyze the differences between states identified by the spiking state model, Bhattacharyya coefficients and distances were computed for pairs of spiking state emission distributions. The Bhattacharyya coefficient between two (potentially non-Gaussian) distributions quantifies their overlap and determines the Bhattacharyya distance *D_BC_* through the relationship *BC* = exp (−*D_BC_*) (56). For each cell *i* and pair of states (*a*, *b*), the state-dependent emission distributions are Poisson distributions of rates *λ_i_*(*a*) and *λ_i_*(*b*), for which the Bhattacharyya distance is

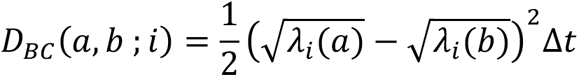

and Δ*t* is the time bin (40 ms). Because we have assumed conditional independence within each state, the full distribution across cells is a product distribution, and the distances add across cells. The Bhattacharyya coefficient between states *a* and *b* is then

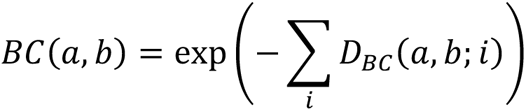

This ranges from 0 to 1, where 1 indicates complete overlap (*λ*_i_(*a*) = *λ*_i_(*b*)) and 0 is no overlap.

#### Statistical tests for cell type differences across recordings

To compare the effects of FS and RS neurons on model fit quality, we compare two metrics. First, we compare the loss in the likelihood relative to the original data in single-neuron-shuffle surrogate datasets. For each single neuron, we generate a surrogate dataset from the reserved test-set population raster, in which spike counts are shuffled in time *for that neuron only*. This surrogate preserves the overall firing rates of all neurons in the population, enabling direct comparison to the original data likelihood. For each neuron, we obtain the loss in the log likelihood per time (*T*, total number of bins) by estimating the likelihood of the new state sequence inferred from the surrogate dataset. The difference between single-neuron shuffle surrogate log likelihood and the original log likelihood is taken as an estimate of how correlated each neuron was with the population state.

The second metric we computed was the coefficient of variation (CV) of the rate parameters (*λ_i_*(*m*)) across states *m* for each neuron *i*. This metric is an estimate of how much the rate for a neuron varied with the learned states; higher CVs indicate that the rates vary more across states. For both statistics, we assess the overall difference between RS and FS neurons across recordings using a hierarchical bootstrap test and report *p*-values and *z*-scores (57).

### Analyses of inferred spiking states

#### Ordering spiking states by LFP states

We ordered spiking states based on whether LFP state H or LFP state L was more likely to be observed during the periods in which that spiking state was inferred (Figure 3). Specifically, we computed P(x|s_i_), where s_i_ is the spiking state and x is the LFP state (either H, if LF/HF < 10, or else L), and sorted states by the ratio: P(H|s_i_)⁄P(L|s_i_).

#### Decoding analysis of LFP-spiking state relationships

Multiple classifiers were fit in Figure 4. For all classifiers, the target response is whether LF/HF > 10. Observations were balanced so that 50% accuracy is chance and divided into test and training sets. The inputs to the different types of classifiers are (i) summed spike count across the population, (ii) the population spike count vector, (iii) the inferred spiking state, or (iv) the single neuron spike counts. The classifier was a linear SVM.

#### Spiking state-dependent analysis of sensory-evoked responses

The evoked response was quantified as the difference between the spike count in the post-stimulus window [5ms, 25ms] and the pre-stimulus window [−20ms, 0ms]. Evoked spike counts were computed for each trial, and we wanted to know if the average evoked response in each state was higher or lower than would be expected from the null hypothesis that evoked responses have no dependence on pre-stimulus spiking state. The number of trials assigned to each state was not identical, thus we determined significance using a shuffle test. Specifically, we randomly assigned trials to states matching the number of trials that were classified to that state and computed the average evoked rate by state in each random draw. This was repeated 200 times for each recording and for each number of spiking states (2 to 10) to generate the shuffled distribution of average evoked response. The empirical mean state-dependent evoked spike count was compared to this distribution. Significance was determined by a two-tailed 1% cut-off, so ‘facilitated’ responses were those in which the empirical mean state-dependent evoked spike count scored above the 99.5% percentile of all shuffled mean evoked responses.

#### Information calculation between whisking activity and spiking state identity

Mutual information quantifies the difference between the joint distribution P(X, Y) of quantities X and Y compared to the distribution P(X)P(Y) in which X and Y are independent, providing a generalized measure of correlation (including non-linear correlation) between quantities X and Y. Specifically, the mutual information (in bits) is 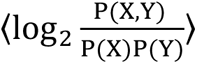, where angle brackets denote an average over the joint distribution P(X, Y). For each calculation of mutual information (Fig. 6), X is the whisking energy, computed in each 40-ms time bin (i) from videography and discretized into four equipartition bins (W_i_ ∈ {0,1,2,3}). To compute information between whisking and spiking states (“Y” is the spiking state), we identified the maximum-likelihood spiking state (s_spk_(i) ∈ {0, 1, … N_S_}) in each time bin and estimated the mutual information between W_i_ and s_spk_(i). To compute information between whisking energy and LFP state (“Y” is LFP state), we assigned LFP state to L or H according to whether r>0 or r<0. Error bars were estimated by resampling over half the data.

#### Code accessibility

Custom MATLAB code to generate figures and perform analyses can be accessed from GitHub following publication.

